# Characterizing phytoplankton communities in the absence of resource-based competition

**DOI:** 10.1101/2022.06.14.496140

**Authors:** Michael J. Behrenfeld, Kelsey M. Bisson, Emmanuel Boss, Peter Gaube, Lee Karp-Boss

## Abstract

Under most natural marine conditions, phytoplankton cells suspended in the water column are too distantly spaced for direct competition for resources to be a routine occurrence. Accordingly, resource-based competitive exclusion should be rare. In contrast, contemporary ecosystem models typically predict an exclusion of larger phytoplankton size classes under low-nutrient conditions, an outcome interpreted as reflecting the competitive advantage of small cells having much higher nutrient ‘affinities’ than larger cells. Here, we develop mechanistically-focused expressions for steady-state, nutrient-limited phytoplankton growth that are consistent with the discrete, distantly-spaced cells of natural populations. These expressions are then encompassed in an ecosystem model that sustains diversity across all size classes over the full range in nutrient concentrations observed in the ocean. In other words, our model does not exhibit resource-based competitive exclusion between size classes. We show that the basis for species exclusions in earlier models is not a reflection of size-dependent nutrient ‘affinities’, but rather a consequence of inappropriate descriptions of non-grazing phytoplankton mortality.

## Introduction

Our interpretation of observed ecological properties derives from our conceptions of the environment experienced by organisms and their interactions with other individuals. These conceptions are inevitably influenced by our own experiences, such that fundamental ecological concepts originally formulated from observations of macro-organisms (birds, mammals, trees, etc.) are often carried forward to very different systems (e.g., microbial communities). For example, principals of resource-based competitive exclusion first deduced from terrestrial communities (Darwin 1927) are commonly assumed equally valid for the phytoplankton (e.g., Hutchinson 1961). At the other end of the spectrum, principals of physical-chemistry can be used to formulate expressions for biochemical reactions, such as the Michaelis-Menten equation for enzymatic reaction kinetics (Michaelis & Menten 1913). The mathematical curve that the Michaelis-Menten equation represents often provides an excellent fit to observational data for far more complex systems (e.g., nutrient uptake in phytoplankton cells), but such results do not imply that associated equation variables carry a similar physiological meaning as those for single enzyme reactions.

The conceptions we have regarding the plankton world are formalized and tested in ecosystem models. For oligotrophic ocean regions, these models often predict the proliferation of small phytoplankton species at the expense (i.e., exclusion) of larger species (e.g., Poulin & Franks 2010, Taniguchi et al. 2014, Dutkiewicz 2020, Acevedo-Trejos et al., 2015). This outcome is interpreted as the consequence of resource-based competitive exclusion, where smaller cells have a greater ‘affinity’ for nutrients and thus can draw nutrients down to a level that will not sustain larger cells. These predictions of size-dependent exclusions are inconsistent with observations. In the oligotrophic ocean, small species may be both numerically- and biomass-dominant, but the phytoplankton size distribution is nevertheless a continuum, where large cells persist at lower abundances (e.g., Venrick 1990, Reynolds & Stramski 2021). This fundamental inconsistency, which has been noted previously (e.g., Ward et al. 2013), motivated the current exploration of how we might better conceive of, and subsequently model, the growth environment experienced by phytoplankton and their interactions with other individuals.

In the narrative below, we begin with a depiction of aquatic ecosystems where individual phytoplankton are distantly spaced across nearly the full range of naturally-occurring cell abundances. In such seascapes, resource-based competitive exclusion is unlikely, which is at odds with the above noted loss of species diversity in plankton ecosystem models under low-nutrient conditions. This contradiction suggests that there is something fundamentally incorrect about the models. One possibility is that the problem lies with the model treatment of phytoplankton taxonomic groups simply as integrated nutrient (e.g., nitrogen) stocks sharing (i.e., competing for) common nutrient resources, rather than from the perspective of how individual cells experience their growth environment. To explore this possibility, we develop a diffusion-based model framework that is consistent with the growth of distantly-spaced, non-competing individuals and which derives from the mechanistic underpinnings of production-resource relationships observed in laboratory populations under light-limiting and nutrient-limiting conditions. A problem that arises from this approach is that the underlying physics yield an untenable initial prediction of extreme size-dependent differences in phytoplankton division rates. However, this issue is resolved when the model is applied to nutrient conditions reflective of natural oceanographic settings. Interestingly, our diffusion-based expression can be equated to a Michaelis-Menten functional form, implying that the apparent problem with ecosystem models does not lie in the treatment of nutrient uptake by integrated stocks. With this finding, we then modify a published multi-species ecosystem model and demonstrate how this revised version sustains all modeled phytoplankton size classes across the full domain of ultra-oligotrophic to eutrophic nutrient conditions. As you will discover at the end of this narrative, the common loss of species diversity in contemporary ecosystem models under low nutrient conditions has nothing to do with resource-based competitive exclusion, so this has been an error in interpretation. We hope you enjoy our story and ‘stick with it’ to the end, as we suspect our findings may be a bit surprising. They certainly were for us.

## Cell spacing and resource competition

Edward Hulburt, using data collected during a series of field campaigns spanning from the north Atlantic gyre to highly productive coastal waters, quantified the average spacing between nutrient depletion zones (i.e., ‘boundary layers’) associated with neighboring phytoplankton cells (Hulburt 1970). His analysis indicated that cell concentrations (of microphytoplankton and large nanophytoplankton) greater than ~10^8^ L^−1^ would be required for cell boundary layers to predominantly overlap and, thus, for direct resource competition to ensue. This requirement exceeded observed cell concentrations by at least two orders of magnitude across all sampled environments, save two shallow coastal estuaries. Accordingly, he concluded that this spatial separation implies that *“no cell can affect any other, that no species can interact with another, through competition for nutrients”* in most natural environments and that this lack of resource-based competition is the explanation for Evelyn Hutchinson’s (1961) ‘Paradox of the Plankton’.

In 1998, David Siegel greatly expanded upon Hulburt’s earlier work, quantitatively evaluating the discreteness of phytoplankton across the cellular size domain. He introduced two distribution variables (*DV*) to assess the likelihood of overlapping ‘spheres of influence’ (i.e., boundary layers). In the spatial domain, *DV_λ_* was defined as the ratio of the diameter of a phytoplankton’s sphere of influence (*d_soi_*) to the average distance between adjacent cells (*λ*):

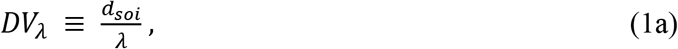

where *DV_λ_* > 1 indicates overlapping boundary layers and competition for resources, whereas *DV_λ_* < 1 indicates that cells are too far apart to feel the effects of their neighbors. The second distribution variable (*DV_τ_*) was introduced to account for interactions between cells over time:

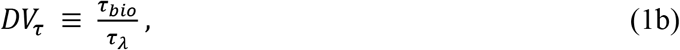

where *τ_bio_* is a characteristic biological time scale (e.g., time between cell divisions) and *τ_λ_* is the time scale that a given phytoplankter will feel the effects of its neighbors. In the case of nutrient competition, *DV_τ_* can be thought of as “*the rate at which neighboring cells are intercepting a given cell’s potential nutrient supply in relation to the cell’s intrinsic nutrient demand over its division cycle”* (Siegel 1998). Here again, *DV_τ_* > 1 indicates that neighboring cells will influence a given phytoplankter’s nutrient supply, while *DV_τ_* < 1 implies minimal competition for nutrients between cells.

To illustrate the relationship between cell size, abundance, and the potential for resource competition, Siegel (1998) assumed spherical cells in a quiescent medium, that a given phytoplankton population is composed entirely of a single cell size, that a cell’s boundary layer is five times larger than the cell’s diameter, and that *τ_bio_* = 1 day [see Siegel (1998) for details and discussion on sensitivity of results to these assumptions]. For these assumptions, the threshold cell abundances for direct resource competition can be found [i.e., solving equations 11 and 13 in Siegel (1998) for *DV_λ_* = 1 and *DV_τ_* = 1] and expressed as a function of cell diameter (Fig. 1a). The salient results here are that threshold abundances increase rapidly with decreasing cell size (note the logarithmic scaling of the y-axis in Fig. 1a) and that across the cell size domain of phytoplakton these thresholds greatly exceed typical cell concentrations found in natural ocean waters.

**Figure 1:**
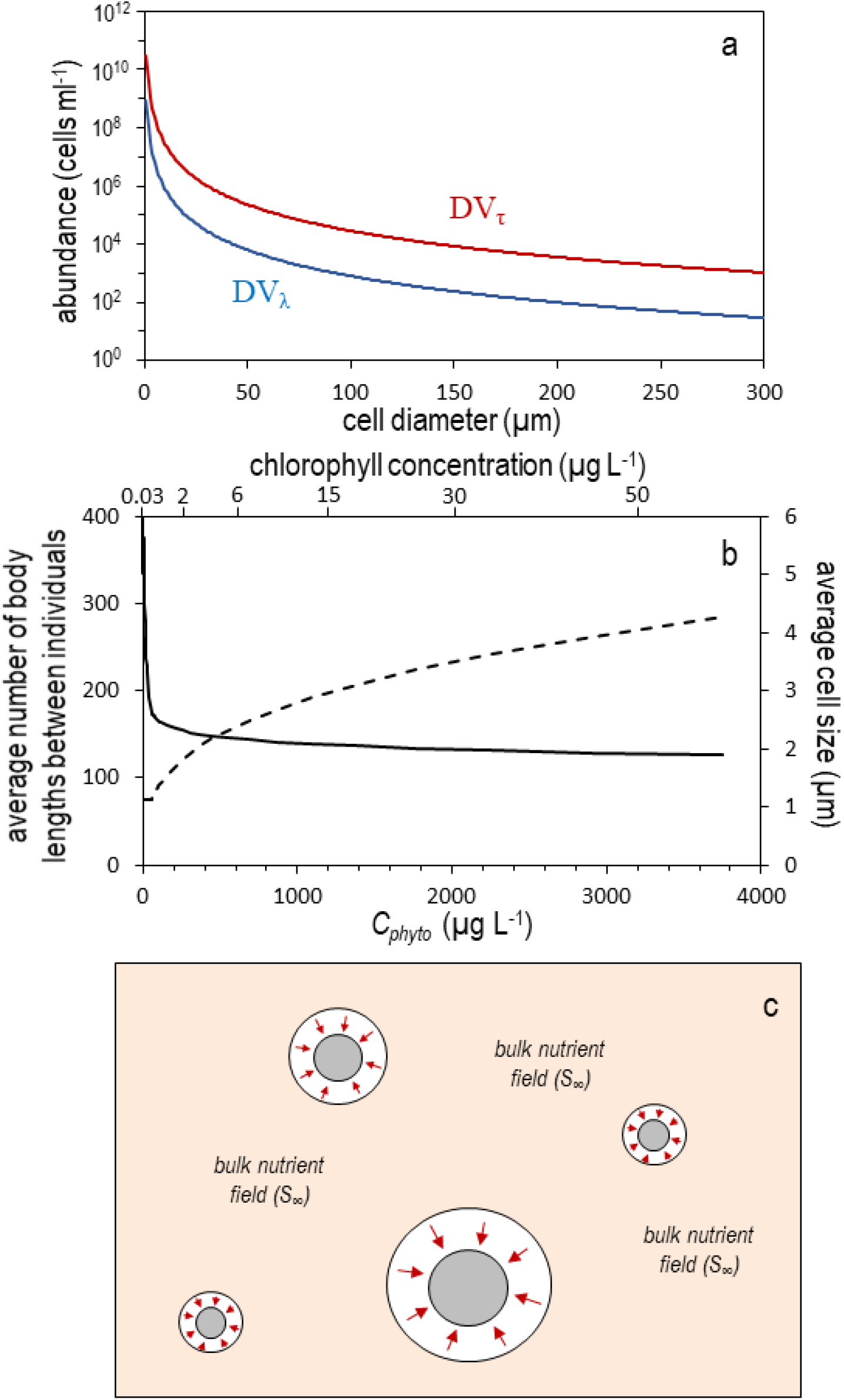
Conceiving the discreteness of phytoplankton communities. (a) Cell abundances for populations of a single cell size required for the spatial (*DV_λ_*) and temporal (*DV_τ_*) distribution variables defined by Siegel (1998) to have a value of one, indicating direct competition for resources is prevalent. Note, these threshold values are notably larger than most natural population abundances. (b) Average number of body lengths between individual phytoplankton cells (left axis, solid line) and average population cell size (right axis, dashed line) for modeled phytoplankton communities with size distributions reflective of natural populations (see text). Cell size is calculated as the cell diameter of the average cell volume. Bottom and top axis give total phytoplankton carbon biomass (*C_phyto_*) and approximate corresponding chlorophyll concentrations. (c) Depiction of phytoplankton in natural waters where cells are distantly spaced and resource acquisition is limited to discrete boundary layers around each cell (outer circles with inward pointing arrows) and has no immediate impact on the far-field resource pool (*S_∞_*) experienced by all cells.

We can further develop our conception, or ‘intuition’, regarding resource competition among phytoplankton by considering populations of a continuous size distribution. In stable lower-latitude oligotrophic systems, phytoplankton abundance is dominated by *Prochlorococcus*, with numbers typically ranging between 2 × 10^4^ and 2 × 10^5^ cells ml^−1^ (Partensky et al. 1999, Martiny et al. 2022) and the phytoplankton community generally exhibits a size distribution with a slope of approximately −4.5 between the logarithm of cell number concentration per unit length and the logarithm of cell diameter (*d*) (Sheldon et al. 1972, Jonasz & Fournier 2011, Huete-Ortega et al. 2012, Marañón 2015, Behrenfeld et al. 2021a). As nutrient stocks in the environment increase, *Prochlorococcus* abundances peak while the abundance of larger cells continues to increase, causing the size distribution slope to tilt upward toward −3 (Behrenfeld et al. 2021a).

For the power-law size spectrum of phytoplankton communities^*^, the total number of individuals [*n* (cells ml^−1^)] between two limits in size [*d_min_, d_max_* (μm)] can be calculated as:

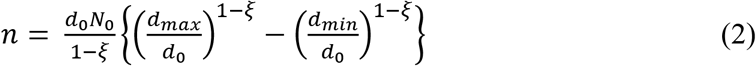

where ξ is the absolute value of the size distribution slope, *d_0_* is a reference cell diameter (μm), and *N_0_* is the particle differential number concentration (ml μm^−1^) at *d_0_*. To demonstrate separation distances between phytoplankton cells in nature, we assign *d_0_* = 1 μm^†^ and assume that ξ = 4.5 as available resources allow *Prochlorococcus* abundances to increase from 2 × 10^4^ to 2 × 10^5^ cells ml^−1^. We further assume that additional increases in nutrients do not affect *Prochlorococcus* concentrations (i.e., remain capped at 2 × 10^5^ cells ml^−1^) but rather result in an increased abundance in larger cells such that ξ decreases from 4.5 to 3.3. With these assumptions, the resultant total number of phytoplankton cells between *d_min_* = 0.6 μm and *d_max_* = 500 μm ranges from 3.2 × 10^4^ to 4.1 × 10^5^ cells ml^−1^ and the average cell size of the population increases from 1.1 to 4.3 μm ξ as decreases from 4.5 to 3.3 (Fig. 1b, dashed line, right axis). Assuming spherical cells and a non-diatom cellular carbon (*C*, pg cell^−1^) to volume (*Vol*, μm^3^) relationship of *C* = 10^−0.665+0.939*Log*_10_(*Vol*)^ (Menden-Deuer & Lessard 2000), the above defined phytoplankton populations span a biomass (*C_phyto_*) range of 6 to ~3700 ng ml^−1^ (lower axis in Fig. 1b; noting here that ng ml^−1^ = μg L^−1^), or a chlorophyll (*Chl*) range of approximately 0.03 to 60 ng ml^−1^ assuming an increase in *Chl:C_phyto_* from 0.006 to 0.02 as biomass increases (due to self-shading). Thus, the modeled phytoplankton populations span a range from highly oligotrophic to highly eutrophic conditions, yet across this full range neighboring cells remain separated on average by 380 to 130 body lengths (Fig. 1b, solid line, left axis).

As a final illustration of the diluteness of phytoplankton in nature, we can calculate the total cell volume per milliter of water (*Vol_phyto_*) as:

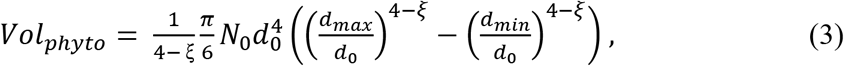

which, for a size range of 0.6 – 500 μm diameter, yields the result that phytoplankton only occupy 0.0000024% to 0.0017% of the volume in which they are suspended for the oligotrophic to eutrophic conditions considered above.

The foregoing analyses provide a general ‘feel’ for the distantly-spaced growth conditions experienced by phytoplankton in a steady-state environment, which is depicted schematically in figure 1c (‘schematically’ because in nature cells are much further apart than illustrated). Of course, this schematic is incomplete and fails to recognize interactions that result from relative movements among cells due, for example, to active swimming, differences in sinking rates, small-scale turbulence, and other processes that bring cells into close proximity (even colliding and forming aggregates). These interactions can result in fleeting competitions for resources. In addition, the abundance of different phytoplankton groups also influences the far-field nutrient concentration (*S_∞_*) experienced by all individuals (much like a change in abundance influences the light field experienced by all cells). For example, during the non-steady-state condition of a phytoplankton bloom, accumulation of a blooming species reduces the value of *S_∞_* experienced by all species (because resource is being drawn from the environment and sequestered into biomass). Likewise, in a nutrient-limited steady-state system, the biomass of different size classes is determined by predator-prey relationships that scale (not necessarily in a 1-to-1 fashion) with division rate. We will return to this latter idea below, but for the moment the central message conveyed in figure 1 and above is that, to first order, phytoplankton experience their world as discrete entities with boundary layers rarely (on a day-to-day basis) overlapping with those of other individuals.

Given the above conception, the question arises whether discreteness of the phytoplankton requires a modification in how we model aquatic ecosystems? Essentially the same question was asked by David Siegel (1998), who was considering the model construct:

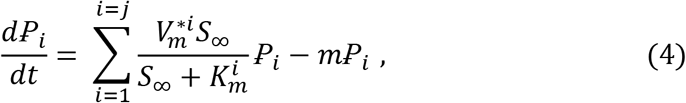

where 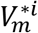 is the maximum specific uptake rate and 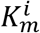 is substrate concentration at 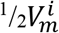 for the *i^th^* phytoplankton group 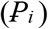 and *m* is the specific loss rate. In this formulation where growth rate is described in a Michaelis-Menten fashion, the phytoplankton community is simply expressed as the integrated stock of some nutrient element without explicit representation of individuals, thus it was concluded that:

> *“Discreteness in phytoplankton…means that formulations of phytoplankton growth and interaction based upon assuming that planktonic organisms are fluid variables* [as in equation 4 above] *are inappropriate for modeling phytoplankton population variations in natural waters. These models assume a priori that phytoplankton populations are distributed continuously, where every cell will uniformly and instantaneously feel the effects of its neighbor”* (Siegel 1998)

The current authors have also voiced a similar concern (Behrenfeld et al. 2021a,b). But, does the omission of discreteness in equation 4 actually imply ‘uniform and instantaneous’ interactions that lead to competitive exclusion? We suggest below that, in fact, it does not. We build toward this conclusion by considering fundamental properties of phytoplankton production-resource relationships common to both light-limited and nutrient-limited growth.

## Production-resource relationships

The relationship between phytoplankton production (e.g., photosynthesis, cell division) and resource supply can be described by equations requiring only two parameters. For the familiar Michaelis-Menten-type expression (e.g., Eq. 4), these parameters are *V_m_* and *K_m_* when describing nutrient-limited growth, with the latter term representing the fore-noted ‘affinity’ for a given resource. This interpretation of *K_m_* has been promoted by laboratory competition experiments where species with lower *K_m_* (i.e., greater affinity) ultimately displace those with higher *K_m_* (e.g., Tilman 1981). However, employing Michaelis-Menten-type expressions in ecosystem models raises a couple important issues. First, the Michaelis-Menten equation (Michaelis & Menten 1913) was developed to describe single substrate-enzyme-product systems and is mechanistically grounded in physical-chemistry. When applied to whole-cell properties, this mechanistic basis for the equation (thus, interpretation of model variables) is compromised. Second, it is not immediately clear whether expressions, such as equation 4, provide robust predictions for populations of distantly-spaced, non-competing individuals, as depicted for phytoplankton in the previous section. In the following, we therefore consider a mechanistic interpretation of production-resource relationships that we propose provides some insight on these apparent challenges with Michaelis-Menten-based formulations. We begin by considering two fundamentally-different light-limited experimental conditions and then draw parallels between these responses and nutrient-limited experimental data to ultimately build a diffusion-focused expression appropriate for describing phytoplankton growth in natural communities.

Let us first consider the familiar experimental procedure of determining ‘Photosynthesis-Irradiance’ relationships (i.e. a ‘*P-I* curves’), whereby samples from a phytoplankton population acclimated to a given growth irradiance (*I_g_*, mol quanta m^−2^ d^−1^) are exposed to a range of light levels (*I_E_*, mol quanta m^−2^ d^−1^) for a sufficiently brief period that physiological acclimations are negligible. Resultant *P-I* curves (e.g., Fig. 2a) are characterized by an initial linear increase in photosynthesis that is defined by photon flux, the number of absorbing (pigment) targets, and the functional absorption cross-section per target. At higher light intensities, the rate of photon capture begins to approach the cell’s capacity to utilize the light-driven production of ATP (adenosine triphosphate) and reductant (nicotinamide adenine dinucleotide phosphate, NADPH). This capacity, which defines the maximum (light-saturated) rate of photosynthesis (*P_m_*), is ultimately determined (primarily) by turnover of the Calvin-Benson-Bassham Cycle (Sukenik et al. 1987, Glover 1989, Fisher et al. 1989, Rivkin 1990, Orellana and Perry 1992, Flynn and Raven 2017). The value of *P_m_* is a species-specific property aligned with its evolutionarily-selected maximum division rate (*μ_m_*). Thus, *P-I* curves are defined by a physical process (photon capture) and a biological limit (*P_m_*) and, logically, traditional equations describing these curves are formulated in these terms, such as (Jassby & Platt 1976):

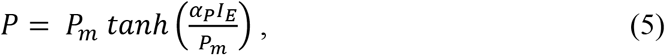

where the slope, *α_P_*, is the photon capture efficiency at low light [m^2^ (mol quanta)^−1^]. *P-I* expressions, such as equation 5, yield relatively abrupt transitions between light-limited and light-saturated photosynthesis and commonly provide a suitable fit to observational data (e.g., Fig. 2a). An ‘emergent property’ of *P-I* curves is the light saturation index, 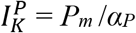. For the hyperbolic tangent model (Eq. 5), *P* = 0.76 *P_m_* when 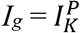.

**Figure 2:**
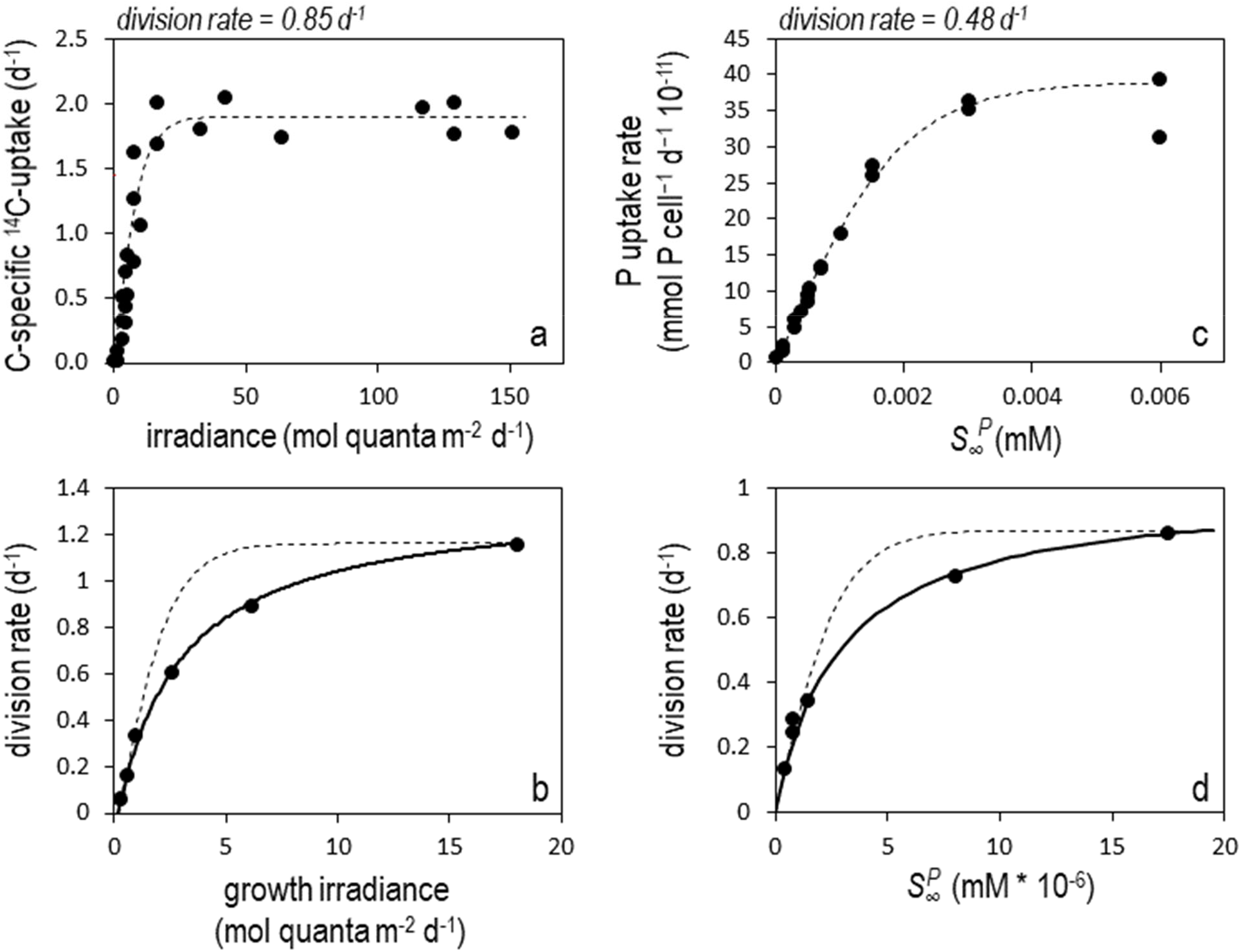
Short-term and acclimated production-resource relationships for light-limited and nutrient-limited phytoplankton populations. (a) Short-term (20 minute) carbon-specific ^14^C uptake as measured by Fisher & Halsey (2016) for *Thalassiosira pseudonana* (Hustedt) Hasle et Heimdal (CCMP 1355) cultures acclimated to a light-limited growth rate of 0.85 d^−1^. Dashed line = fit of equation 5. (b) Cell division rates observed by Laws & Bannister (1980) for *Thalassiosira weissflogii* (previously, *Thalassiosira fluviatilis*) acclimated to a range in growth irradiance (*I_g_*, x-axis). Solid line = fit of equation 6. Dashed line = application of equation 5. (c) Short-term (8 minute) PO_4_ uptake (atto-mol = 10^−15^ mmol) measured by Laws et al. (2011b) for *Pavlova lutheri* (Droop) J.C. Green maintained in chemostats at a PO_4_-limited growth rate of 0.48 d^−1^ and then rapidly exposed to a range of concentrations (x-axis). Dashed line = fit of equation 7. (d) Cell division rates observed by Laws et al. (2011a) for *Tetraselmis suecica* (Kylin) Butcher in steady-state PO_4_-limited chemostat cultures. Solid line = fit of equation 8. Dashed line = application of equation 7. (c,d) x-axis = measured far-field PO_4_ concentration 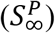.

In a similar manner to a *P-I* curve, cell division rates (*μ*, d^−1^) can be measured across populations acclimated to different growth irradiances (i.e., a ‘*μ-I_g_’* curve). Emergent *μ-I_g_* curves are again saturating functions of light level (Fig. 2b), but these relationships differ significantly from *P-I* curves because each population has had sufficient time to optimize its physiology according to its growth conditions. This optimization involves the tuning of a plethora of cellular properties, including investments in photosynthetic machinery, nutrient uptake systems, and respiratory pathways. While generalized models of this acclimation process have been developed (e.g., Geider 1998), we currently do not have adequate knowledge on evolutionary histories and life strategies to *a priori* predict the unique ‘solutions’ expressed by different species. What we can say is that observed *μ-I_g_* curves are again well described (solid black line Fig. 2b) as functions of the physical process of photon capture efficiency at low light [*α_μ_*, m^2^ (mol quanta)^−1^] and a species-specific biological limit on *μ_m_* (d^−1^) at high light. However, *μ-I_g_* relationships generally follow a rectangular hyperbolic form that can be expressed as (modified from Thornley 1976):

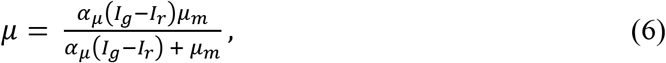

where *I_r_* (mol quanta m^−2^ d^−1^) is the light level at which primary production equals the maintenance respiration rate^‡^. The significant influence of cellular optimization on the *μ-I_g_* relationship is reflected in Fig.2b by the poorness of fit when equation 5 is applied to the data (dashed line). For *μ-I_g_* data, an ‘emergent’ light saturation index can again be defined as 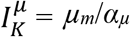, but in the case of the rectangular hyperbolic (Eq. 6), *μ* = 0.5 *μ_m_* when 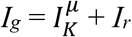.

Production-resource relationships for nutrient-limited populations exhibit very similar properties as light-limited systems. For example, Laws et al (2011a) exposed cells from a steady-state phosphate-limited population of the haptophyte, *Pavlova lutheri*, growing at *μ* = 0.48 d^−1^ to a range of phosphate concentrations 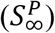 and measured short-term (8 minute) uptake rates (*v*, mmol cell^−1^ d^−1^). Analogous to equation 5, the observed uptake-substrate (*v-S_∞_*) curve is well described by (Fig. 2c):

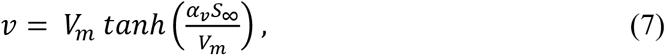

where α_*v*_ (L d^−1^ cell^−1^) is an initial slope (akin to α_P_) defined by the physical flux of nutrients to the cell surface (i.e., diffusion through the cells boundary layer) and capture by membrane transporters (the ‘targets’), while *V_m_* is the nutrient-saturated maximum uptake rate (mmol cell^−1^ d^−1^). As noted by Laws et al (2011a) and later by Flynn et al. (2018) for nitrogen-limited cultures of *Emiliania huxleyi* and *Heterosigma carterae, v-S_∞_* data are often not well fitted by a rectangular hyperbolic function. Because such data are collected on time scales too short for physiological acclimations, *v-S_∞_* curves reflect cell uptake capacities for a fixed population of membrane transporters and, thus, can be mechanistically described by so-called ‘diffusion-porter’ models (Pasciak & Gavis 1974, Shaw et al. 2014, Armstrong 2008, Aksnes & Cao 2011, Flynn et al. 2018).

Modeling steady-state nutrient-limited phytoplankton growth, on the other hand, requires a description of performance as a function of resource availability (*S_∞_*) akin to a *μ-I_g_* relationship. Here again, optimization of growth results from tuning a plethora of cellular processes (not simply those associated with nutrient uptake and assimilation) and observed relationships reflect evolved ‘solutions’ specific to each species that, again, we currently do not have sufficient mechanistic understanding of to accurately predict *a priori*. Nevertheless, *μ-S_∞_* relationships commonly follow a saturating rectangular hyperbolic form^§^ (e.g., Fig. 2d):

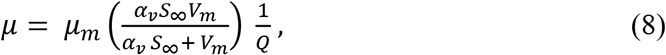

where *α_v_* (L d^−1^ cell^−1^) is, in this case, the slope of uptake versus substrate concentration (mM) for populations acclimated to low nutrient levels and *Q* is the cellular requirement for limiting nutrient (mmol cell^−1^) (note the ratio 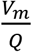 is equivalent to 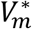 in Eq. 4). As in figure 2b, equation 7 does not provide a suitable fit to *μ-S_∞_* data (dashed line in Fig. 2d), again illustrating the significant influence of cellular optimization in acclimated populations.

A take-home message of the foregoing discussion is that short-term and acclimated production-resource relationships for both light-limited and nutrient-limited conditions can be logically described as functions of a largely physically-defined flux-capture process (i.e., the *α* terms) and an evolutionarily selected for, species-specific maximum division rate or related biological property (i.e., *V_m_. P_m_*). However, the shapes of short-term and acclimated production-resource relationships differ because of the many physiological ‘knobs’ cells can turn in the process of optimization. The outcome of this optimization for nutrient-limiting conditions is a production-resource response that typically follows a rectangular hyperbolic form where the saturation index, *V_m_/α_v_*, corresponds to the nutrient concentration where 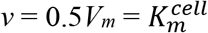. Accordingly, the right hand side of equation 8 can be reorganized by multiplying the numerator and denominator by 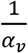 and substituting 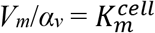 to give:

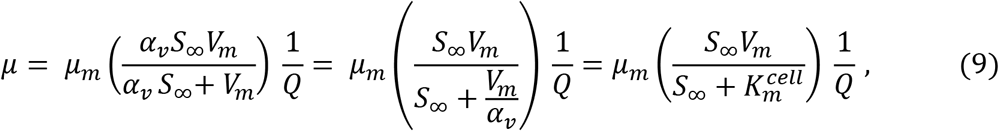

where the bracketed term of the right-most expression is now the Michaelis-Menten form commonly employed in contemporary plankton ecosystem models (e.g., Eq. 4). Here, we have used the term, 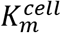, to acknowledge that this evolved property of optimized whole-cell physiology should not be mechanistically thought of as equivalent to the *K_m_* of a Michaelis-Menten single enzyme-substrate-product reaction.

We propose that, when thinking about nutrient-limited phytoplankton growth in nature, equation 8 has a distinct advantage over its converted Michaelis-Menten form in equation 9 (and similarly Eq. 4), despite their mathematical equivalence. Specifically, the Michaelis-Menten form is often interpreted as specifying competitive advantages/disadvantages between phytoplankton species because *V_m_* and 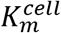 are both viewed as species-specific physiological ‘traits’. While this certainly is the case for *V_m_* and a high *V_m_* can bestow an advantage under elevated resource conditions, the limit on 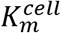 is primarily dictated by the physical process of diffusion. As such, the common assignment of 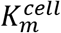 as a metric of nutrient ‘affinity’ [‘*an attractive force between substances or particles’* (Merriam-Webster, https://www.merriam-webster.com/dictionary/)] is misleading, as cells cannot ‘attract’ resources beyond diffusional limits [for a given morphology, relative motion, and assuming equal efficiency in capturing nutrients at the rate they arrive at the cell surface (see below)]. When individuals are as distantly spaced as in natural populations (Fig. 1), their influence on the nutrient field is constrained to their respective boundary layers (outer circles and red arrows in Fig. 1c) and has no immediate impact on the far-field (*S_∞_*) experienced by neighboring cells. So long as the phytoplankton of such a community are consumed and their nutrients recycled at a rate equivalent to population uptake (i.e., steady-state conditions), *S_∞_* remains unchanged and there is, on average, no direct competition between cells, nor does the higher uptake per unit cell volume of smaller phytoplankton directly diminish the uptake and growth of larger cells (we use the term ‘directly’ here because the standing stock of a given phytoplankton type has an indirect influence on other phytoplankton by influencing *S_∞_*, as we previously discussed). The key point is that, in nature, a species with a low 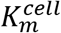 does not have an ability to draw down *S_∞_* such that it can competitively exclude other species with higher 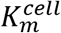. However, this is generally not the case in laboratory competition experiments (e.g., Tilman 1981) where high cell concentrations can result in direct competition and subsequent exclusion (Siegel 1998).

In the previous section, we painted a picture of the phytoplankton world highlighting the discreteness of individual cells. We then asked whether traditional formulations for phytoplankton growth (Eq. 4) found in contemporary ecosystems models are flawed because they treat phytoplankton populations as continuous fields of an elemental stock, rather than as discrete entities. In the current section, we discuss the basis of observed production-resource relationships and provide an expression (Eq. 8) for nutrient-limited growth of distantly-spaced, non-competing phytoplankton. For a population of equally-sized cells, this equation yields the same prediction for steady-state biomass when implemented at the individual level and then integrated over the population as when implemented at the integrated population level. For a population of polydispersed cells, the outcome of equation 8 is likewise the same for these two implementation approaches so long as size-dependences in diffusion are accounted for. As equation 8 is mathematically equivalent to a Michaelis-Menten form, *our conclusion is that there is nothing fundamentally incorrect about applying relationships such as equation 4 when modeling distantly spaced phytoplankton*. Ironically, what may be missing from such equations is, instead, a term accounting for competition under conditions when cells are in close proximity.

If the fore-stated conclusion is valid, then where have previous interpretations gone wrong? We propose that the answer to this question is two-fold. First and as noted above, the thought of 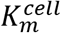 as a species-specific ‘affinity’ acting to deplete *S_∞_* is incorrect and should be replaced by a view that 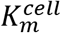 is an ‘emergent property’ of size-dependent diffusion processes, species-specific *μ_m_*, and evolved optimization strategies. Second, previous interpretations of equation 4 are incorrect; specifically, the conclusion that treating phytoplankton biomass as an integrated elemental stock is equivalent to a continuously distributed fluid variable. The fact is that there is nothing about equation 4 that explicitly states how biomass is spatially distributed, only that it has an integrated mass. If information on the size of cells within this mass is retained, then appropriate diffusion rates can be applied in calculating growth rates, irrespective of the spacing between individuals.

The above insights help reconcile earlier conceived issues, but they also make the original problem motivating this study even more vexing. If current expressions in ecosystem models are consistent with growth in a competition-neutral resource landscape (Behrenfeld et al. 2021a,b), then why do these models yield extinctions of most phytoplankton size groups under oligotrophic conditions? As a first step toward answering this question, we will now explore the size distribution of phytoplankton division rates from a diffusion-focused perspective.

## Diffusion-supported phytoplankton division rates

The diffusional flux of nutrients to the surface of a phytoplankton cell is dependent on a variety of factors, including cell size and shape, movement (e.g., sinking, swimming) relative to the surround medium, and the efficiency with which transporter proteins translocate nutrients across the cell membrane relative to the diffusive rate at which they arrive at the membrane [this relative rate determines the concentration gradient between the cell surface (*S_0_*) and *S_∞_*]. Here, we will forgo a detailed description of molecular diffusion and, instead, refer interested readers to the rich literature that already exists describing solute flux across cell boundary layers (e.g., Munk & Riley 1952, Jumars et al. 1993, Karp-Boss et al. 1996, Berg 2018, Kiørboe 2018). For simplicity, we will assume that phytoplankton are spherical cells, such that the diffusional flux ((*F_D_*) to the cell surface in the absence of relative motion is described by:

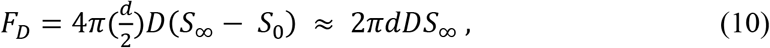

where *d* is cell diameter, *D* is the diffusion coefficient for a given nutrient type, and the right-most expression assumes that the cell is a perfect absorber (i.e., *S_0_* = 0). Equation 10 states that the nutrient flux to a phytoplankton scales with cell diameter. Accordingly, the cell-volume-specific flux 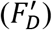 for a spherical cell is:

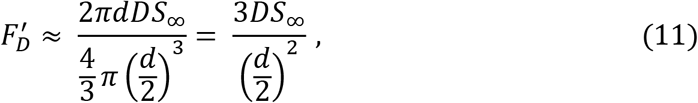

which predicts that, if growth rate is limited purely by diffusion, the size-distribution of division rates will not scale in proportion to the surface:volume ratio (i.e., 1/*d*; upper heavy black lines in Fig. 3a,d) but rather with the inverse square of cell diameter (i.e., 1/*d*^2^; lower heavy black lines in Fig. 3a,d). To place this initial prediction in context, *it implies that a nutrient-limited 1 μm cell will be dividing 10,000 times faster than a nutrient-limited 100 μm cell*. Clearly, this prediction is inconsistent with reasonable size-dependent changes in *μ* for natural populations (Laws 2013). Nevertheless and as noted by Jumars et al. (1993), even large cells are limited in their ability to alleviate this strong constraint of diffusion through their relative motions or morphological adaptations (see also Karp-Boss et al. 1996).

**Figure 3:**
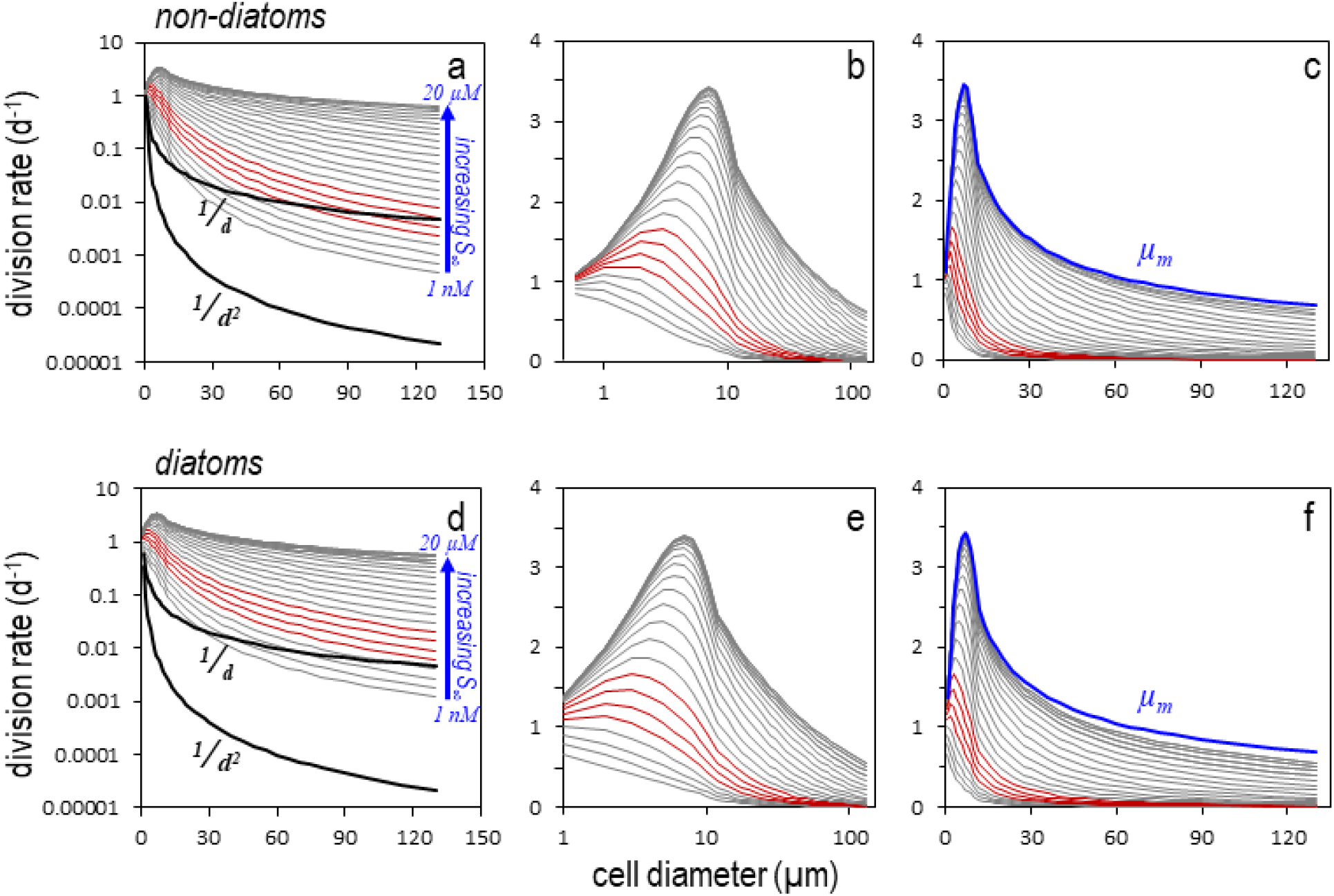
Diffusion-supported phytoplankton division rates as a function of cell size predicted for a range in far-field nutrient concentrations (*S_∞_*) reflective of highly oligotrophic to highly eutrophic natural waters. (a-c) Non-diatoms. (d-f) Diatoms. (a,d) Lower heavy black line = initial prediction for diffusion-limited growth at all cell sizes. Upper heavy black line = division rate prediction if following cellular surface:volume ratios. Grey lines = size-dependent division rates for *S_∞_* ranging from 1 nM to 3 μM (blue labeling). Red lines = division rates for typical *S_∞_* of biologically-available nitrogen in oligotrophic ocean gyres. (b,e) Same data as in left column but with normal y-axis and log-transformed x-axis to better reveal size-dependent division rates of small cells. (c,f) Same data as in left column but with normal axes. Blue line = envelope in size-dependent maximum division rates (*μ_m_*) from Behrenfeld et al. (2021c).

Resolution of the above issue emerges by combining estimates of diffusional flux (Eq. 10) with a *μ-S_∞_* relationship (Eq. 8) for nitrogen-limited growth. In oligotrophic ocean regions, surface layer ammonium (NH4) concentrations commonly range from <3 nM (i.e., detection limit) to 10’s of nM, while summed nitrate and nitrite levels range from undetectable to slightly less than 10 nM (Brzezinski 1988, Jones 1991, Raimbault et al. 2008, Ellwood et al. 2013, Hashihama et al. 2015, https://hahana.soest.hawaii.edu/hot/hot-dogs/interface.html). In more productive ocean regions, available nitrogen sources can exceed μm concentrations (Voss et al. 2013). We therefore consider here far-field nitrogen concentrations of 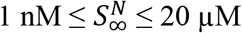.

Cell division rates over our range in 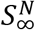 were determined for diatoms of diameter 1 to 130 μm and for other phytoplankton over the size range 0.6 to 130 μm. The diffusional flux of nutrient was calculated from equation 10 assuming *S_0_* = 0 and a constant *D* = 1500 μm^2^ s^−1^ (i.e., ignoring, for example, effects of temperature). The influence of relative motion on *F_D_* was estimated by first calculating a size-dependent characteristic velocity (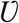; μm s^−1^) for swimming by non-diatoms 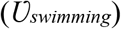 following (Sommer 1988):^**^

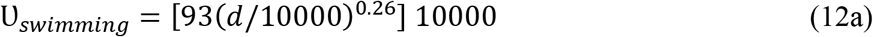

and for sinking 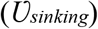 in diatoms of *d* ≥ 8 μm (i.e., 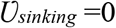 for *d*< 8 μm) based on data from Waite et al. (1997; their figure 7a):

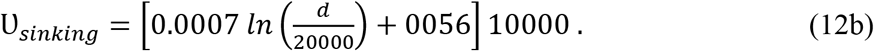

Péclet numbers (*P_e_*) for both phytoplankton groups were then calculated as:

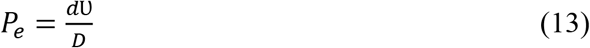

and used to assess Sherwood numbers (*Sh*) following (Karp-Boss et al. 1996):

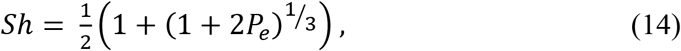

where *Sh* quantifies the enhancement in diffusive flux relative to *F_D_* of a non-moving cell (Eq. 10). Finally, Jumars et al. (1993) evaluated the relationship between membrane transporter abundance and the fraction of *F_D_* captured by a cell. This analysis revealed a remarkable efficiency (also see Berg & Purcell 1977), such that less than 0.1% of a membrane needs to be occupied by transporters to collect ~50% of the diffusive flux. Here, we will assume a 90% capture efficiency, which would correspond to ~1% membrane coverage (Jumars et al. 1993; their figure 3). The *potential* nutrient flux available for assimilation (*A_F_*) is thus:

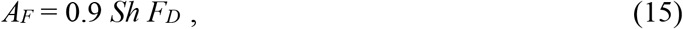

which can be substituted in equation 8 as 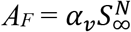.

To calculate division rate (Eq. 8), a size dependent estimate of *V_m_* is needed. In an earlier study (Behrenfeld et al. 2021c), we found that measured maximum division rates (cell doublings per day) reported in the literature fall within an upper envelope that increases with cell size up to *d* ~ 7 μm and then decreases with size in a manner following a power function at *d* ≥ 15 μm. This envelope is described by (upper blue curves in Fig. 3c,f):

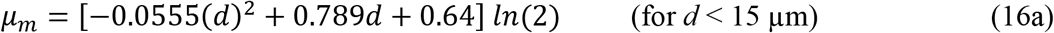

and

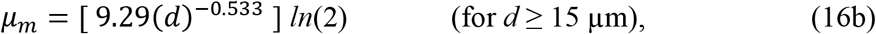

where multiplication by *ln*(2) in each equation coverts maximum doublings per day (reported in Behrenfeld et al. 2021c) to maximum specific division rates (*μ_m_*, d^−1^). As *V_m_* (fg N cell^−1^ d^−1^) is the rate of nutrient uptake (in this case nitrogen) required to support *μ_m_*, it can be estimated as the product of cellular carbon and the nitrogen:carbon ratio at *μ_m_* (*N:C_m_*):

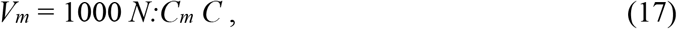

where *C* (pg C cell^−1^) was calculated following Menden-Deuer & Lessard (2000) as stated above for non-diatoms and as *C* = 10^−0.541+0.811*Log*_10_(*Vol*)^ for diatoms, *N:C_m_* was estimated from *N:C* = 0.0762*μ* + 0.0389 based on Laws & Bannister (1980; their Table 1 for nutrient-limited cultures), and the scalar, 1000, converts *C* in pg C cell^−1^ to fg C cell^−1^. Application of equations 15 and 17 in the rectangular hyperbolic element of equation 8 yields the nutrient assimilation rate for a given cell size and 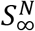. Relating this rate to *μ* requires accounting for growth-rate-dependent changes in nutrient requirements specified by our equation for *N:C*. Realized division rates for each value of 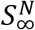 were thus determined by first calculating the division rate that is supported by a given assimilation rate, assuming *N:C* = *N:C_m_*, and then iteratively adjusting *N:C* and *μ* from this initial estimate until stable values were achieved. For all combinations of cell size and 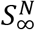, this stabilization was achieved within 25 iterations.

**Table 1.** **S**ymbols/abbreviations, definitions, and units (in order of appearance in manuscript).

| Symbol/<br>abbrev. | Definition | Units |
| --- | --- | --- |
| $\lambda$ | average distance between individual cells | $\mu\text{m}$ |
| $d_{soi}$ | diameter of cell boundary layer | $\mu\text{m}$ |
| $\tau_\lambda$ | interaction time scale between cells | s |
| $\tau_{bio}$ | characteristic biological time scale | s |
| $n$ | number of cells per unit water volume | cells $\text{m}^{-3}$ |
| $d$ | cell diameter | $\mu\text{m}$ |
| $d_0$ | reference diameter | $\mu\text{m}$ |
| $N_0$ | particle differential number concentration at $d_0$ | $\text{ml } \mu\text{m}^{-1}$ |
| $\xi$ | absolute value of the size distribution slope | unitless |
| $Vol$ | cell volume | $\mu\text{m}^3$ |
| $C$ | phytoplankton cellular carbon content | $\text{pg cell}^{-1}$ |
| $C_{phyto}$ | carbon mass of phytoplankton population | $\text{ng C ml}^{-1}$ |
| $Chl$ | chlorophyll mass of phytoplankton population | $\text{ng Chl ml}^{-1}$ |
| $Vol_{phyto}$ | total cell volume per unit water volume | $\text{ml ml}^{-1}$ |
| $S_\infty$ | far-field nutrient concentration | $\text{mmol m}^{-3}$ or mM |
| $P$ | phytoplankton biomass in nutrient units | $\text{mmol m}^{-3}$ |
| $V_m^*$ | maximum specific nutrient uptake rate | $\text{d}^{-1}$ |
| $K_m$ | Michaelis-Menten half-saturation substrate concentration | $\text{mmol m}^{-3}$ |
| $m$ | linear specific loss rate of phytoplankton | $\text{d}^{-1}$ |
| $I_g$ | Light level to which phytoplankton are acclimated | $\text{mol quanta m}^{-2} \text{d}^{-1}$ |
| $P$ | carbon –specific photosynthetic rate | $\text{d}^{-1}$ |
| $P_{max}$ | carbon –specific light-saturated photosynthetic rate | $\text{d}^{-1}$ |
| $\mu_m$ | maximum specific rate of cell division | $\text{d}^{-1}$ |
| $\alpha_P$ | light-limited slope of $P$-$I$ relationship | $\text{m}^2 (\text{mol quanta})^{-1}$ |
| $I_K^P$ | light-saturation index of $P$-$I$ curve | $\text{mol quanta m}^{-2} \text{d}^{-1}$ |
| $\mu$ | specific rate of cell division | $\text{d}^{-1}$ |
| $\alpha_\mu$ | light-limited slope of division rate-irradiance (μ-$I_g$) relationship | $\mu\text{mol C m}^2 (\mu\text{mol quanta})^{-1}$ |
| $I_r$ | light level where primary production equals the maintenance respiration rate | $\text{d}^{-1}$ |
| $I_K^\mu$ | light-saturation index of μ-$I_g$ relationship | $\mu\text{mol quanta m}^{-2} \text{d}^{-1}$ |
| $v$ | nutrient uptake rate | $\text{mmol cell}^{-1} \text{d}^{-1}$ |
| $V_m$ | maximum nutrient uptake rate | $\text{mmol d}^{-1}$ |
| $\alpha_v$ | initial slope of the relationship between $v$ and $S_\infty$ | $\text{L d}^{-1}$ |
| $Q$ | cellular requirement for limiting nutrient | $\text{mmol cell}^{-1}$ |
| $K_m^{cell}$ | nutrient concentration where $v = 0.5 V_m$ for cells acclimated to different $S_\infty$ | mM |
| $S_0$ | concentration of given nutrient at the cell surface | $\text{mmol m}^{-3}$ or mM |
| $F_D$ | diffusional flux of nutrient to a stationary spherical cell | $\text{mmol } \mu\text{m}^3 \text{s}^{-1}$ or $\text{fg } \mu\text{m}^3 \text{s}^{-1}$ |
| $D$ | diffusion coefficient | $\mu\text{m}^2 \text{s}^{-1}$ |
| $F_D'$ | cell volume-specific diffusional flux of nutrient to a stationary spherical cell | $\text{mmol } \mu\text{m s}^{-1}$ or $\text{fg } \mu\text{m s}^{-1}$ |
| $U$ | characteristic cell sinking or swimming velocity | $\mu\text{m s}^{-1}$ |
| $Pe$ | Péclet number | unitless |
| $Sh$ | Sherwood number | unitless |
| $A_F$ | Potential nutrient flux available for assimilation assuming 90% capture efficiency and accounting for cell movement | mmol $\mu\text{m s}^{-1}$ or fg $\mu\text{m s}^{-1}$ |
| $P_i^N$ | total nitrogen inventory of the $i^{\text{th}}$ phytoplankton size class | mmol $\text{m}^{-3}$ |
| $Z_i^N$ | total nitrogen inventory of the $i^{\text{th}}$ zooplankton size class | mmol $\text{m}^{-3}$ |
| $f_i$ | feeding size range of grazers and carnivores with respect to mean prey size | unitless |
| $g_1$ | zooplankton grazing rate | $\text{m}^3 \text{mmol}^{-1} \text{d}^{-1}$ |
| $g_2$ | zooplankton ingestion efficiency | unitless |
| $g_3$ | zooplankton linear mortality rate | $\text{d}^{-1}$ |
| $g_4$ | zooplankton density-dependent mortality rate | $\text{m}^3 \text{mmol}^{-1} \text{d}^{-1}$ |
| $\kappa$ | media outflow rate in Ward et al. (2013) chemostat model | $\text{d}^{-1}$ |

The outcome of the above formulations is shown in figure 3 for non-diatoms (top) and diatoms (bottom), where the left panels are plotted with a log-transformed y-axis, the center panels with a log-transformed x-axis, the right panels with normal axes, and in all panels typical oligotrophic conditions of 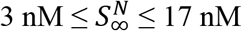 are indicated by red lines. A key finding here is that size spectra for phytoplankton division rates fall far from a scaling with the square of cell diameter (1/*d*^2^), as initially predicted, when based on diffusion for realistic 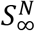 and they even lie above a surface:volume dependence (1/*d*) for all but the lowest 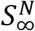. The reason for this is that the diffusional potential (*A_F_*) for small cells is close to or exceeds that necessary to support *μ_m_* even at very low 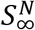, such that any residual changes in *μ* with increasing nutrient supply simply reflect the slowly-saturating nature of evolutionarily-optimized *μ-S_∞_* relationships (e.g., Fig. 2d). Because the steady-state biomass of a given phytoplankton size class is determined by *μ*-dependent predator-prey relationships, expectations from these results when applied to an ecosystem model are an even further dampening in the range of equilibrium biomasses across size classes (compared to the range in size-dependent *μ*) at low nutrients and a much stronger potential for proliferation of large species with increasing 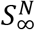, which will tilt the size distribution slope upward toward −3 (Behrenfeld et al. 2021a).

In the next section, we will apply our diffusion-focused approach in a modified version of a published ecosystem model, but before proceeding we conclude the current section with a comparison of our modeled *μ* values and observational data from two chemostat-based studies where measurements of *S_∞_* were reported (Laws et al. 2011a,b) to evaluate if our formulation provides reasonable predictions. In the first of these studies, phosphate-limited cultures of the temperate chlorophyte, *Tetraselmis suecica*, were maintained at steady-state division rates of ~0.16 to ~0.75 d^−1^, which corresponded to 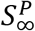 concentrations ranging from ~0.7 to ~5.6 nM, respectively. Applying our above-described diffusion-based approach for an average *T. suecica* cell diameter of 12 μm, a cellular elemental nitrogen:phosphate ratio (*N:P*) of 16:1, and a *μ_m_* of 1.2 d^−1^ (Laws et al. 2011a) yields estimates of *μ* that are highly consistent (R^2^ = 0.96) with observed values (Fig. 4a). This finding implies an optimization strategy in *T. suecica* aimed at fully utilizing the diffusional flux of limiting nutrient to the cell surface. In the second study (Laws et al. 2011b), phosphate-limited cultures of the temperate haptophyte, *Pavlova lutheri*, were maintained at steady-state division rates ranging from ~0.13 to ~0.85 d^−1^, with corresponding 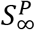 values of ~0.4 to ~17.5 nM, respectively. In this case, application of our diffusion-based approach for an average *P. lutheri* cell diameter of 6 μm, a *N:P* of 16:1, and a *μ_m_* of 1.0 d^−1^ (Laws et al. 2011b) yields correlated (R^2^ = 0.97) but somewhat overestimated values of *μ* compared to observations (Fig. 4b) [note that these same data are very well fitted by an empirically-parameterized version of equation 8 (Fig. 2d)]. Perhaps the observed modest departure from pure diffusion limitation observed in *P. lutheri* implies that the evolved life strategy of this species involves other tactics for success aside from maximizing nutrient utilization. In this regard, it might be noted that the cultured isolate, *P. lutheri* (Droop) Green, was originally obtained from an intertidal location of the Clyde Sea where one might speculate that selection pressures may have been weak for success under low nutrient conditions and perhaps more oriented toward defense (grazer or other) strategies.

**Figure 4:**
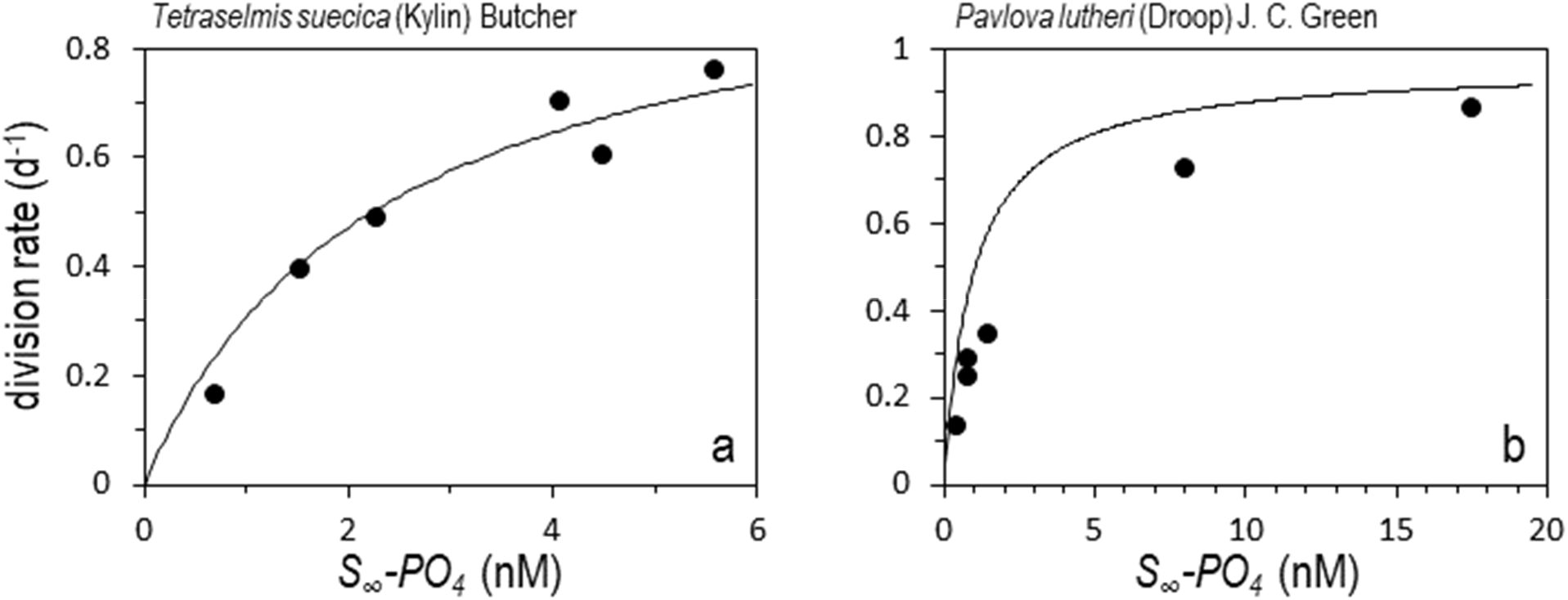
Comparison of model-predicted phytoplankton division rates with measure steady-state rates in PO_4_-limited chemostat cultures. (a) Circles = cell division rates observed by Laws et al. (2011a) for *Tetraselmis suecica* (Kylin) Butcher. Solid line = predicted division rates assuming an average cell size of 12 μm and a maximum division rate (*μ_m_*) of 1.19 d^−1^ (Laws et al. 2011a). (b) Circles = cell division rates observed by Laws et al. (2011b) for *Pavlova lutheri* (Droop) Green. Solid line = predicted division rates assuming an average cell size of 6 μm and a *μ_m_* of 0.98 d^−1^ (Laws et al. 2011b).

## One-dimensional Ecosystem Model

Ecosystem modeling can be a bit of an ‘art form’ in terms of tuning parameters to achieve stable and reasonable results (i.e., when compared to observations) (Siegel 1998, Hilborn & Mangel 2013) and as the complexity of a model increases, understanding its behavior becomes more difficult (Evans & Parslow 1985, Moore 2022). The intention of the following exercise is to answer our initial question regarding why contemporary ecosystem models often yield significant extinctions of larger phytoplankton species under steady-state low-nutrient concentrations, whereas field observations indicate the coexistence of a continuum in phytoplankton sizes. Accordingly, we have chosen here to employ the following minimal equation set such that a clear answer to this question emerges:

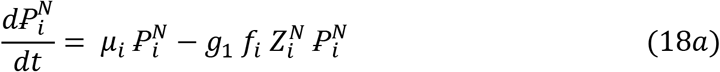

and

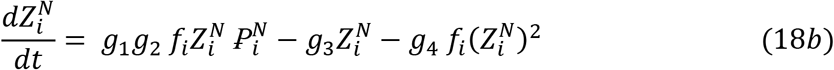

where 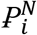 is the total nitrogen inventory (mmol m^−3^) of the *i^th^* phytoplankton size class, phytoplankton mortality is solely due to grazing, 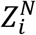 is the nitrogen inventory of the zooplankton population (mmol m^−3^) grazing upon the *i^th^* phytoplankton size class, *μ_i_* is the diffusion-supported division rate (d^−1^) for a given 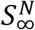 (calculated as described in the previous section), and *f_i_* is defined below. Parameters *g_1_* through *g_4_* represent zooplankton grazing rate, ingestion efficiency, linear mortality rate, and density-dependent mortality rate, respectively, and are assigned size-independent values of *g_1_* = 3.24 m^3^ mmol^−1^ d^−1^, *g_2_* = 0.5 (unitless), *g_3_* = 0.06 d^−1^, and *g_4_* = 1.6 m^3^ mmol^−1^ d^−1^ (Behrenfeld & Boss 2014)

The size structuring of our modeled ecosystem generally follows that described for the *“idealized food-chain model”* developed by Ward et al. (2013). In our ‘baseline’ model for either non-diatoms or diatoms, we include 25 phytoplankton size-classes with diameters ranging from 0.6 μm to 135 μm, where cells in a given size class are 1.25 times larger than those in the class one size smaller. The model also includes 25 zooplankton size classes. We assume that predators of phytoplankton and zooplankton consume prey over a feeding size range that is proportional to (*f_i_* in Eq. 18a,b) their mean prey size (e.g., in the current case we define *f_i_* = 0.5, meaning that a predator with a mean prey size of 100 μm will have a feeding range of 50 – 150 μm, while a predator with a mean prey size of 2 μm will have a feeding range of 1 – 3 μm) (Behrenfeld et al. 2021a, Ward et al. 2012, 2013). Because our focus is on phytoplankton size composition under steady-state (in this case, nitrogen-limited) growth conditions, we do not consider the role of environmental variability. For each model run, 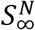 is held constant at a value between 1 nM and 20 μM (stepped every 7 nM between 1 and 35 nM and every 200 nM thereafter) and light levels are assumed constant and saturating for growth. Because 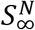 is held constant, phytoplankton nitrogen consumed but not assimilated by grazers and grazer nitrogen lost to linear and density-dependent predation are removed from the modeled system and not recycled. Model runs for each 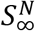 are initiated with 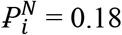 mmol N m^−3^ and 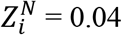 mmmol N m^−3^ for all size classes and then executed for 3 years at ~15 minute time-steps (noting that the 3-year time frame was conservative as actual times for all phytoplankton size classes to reach equilibrium was only 30 days to ~1.5 years, with the longer times required for lower values of 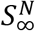). Size diversity for the steady-state populations was assessed as the number of ‘species’ (i.e., size-classes) remaining that contributed at least 0.0001% to total phytoplankton biomass.^††^

The first and most important result from our ecosystem model is that all phytoplankton size classes are retained under all values of 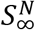 for both our non-diatom and diatom communities (Fig. 5a, red & yellow line). This finding contrasts starkly with results from earlier ecosystem models where only the smallest species persist at low 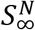 (e.g., Poulin & Franks 2010, Taniguchi et al. 2014, Dutkiewicz et al. 2020). This difference may seem rather surprising given our earlier statement that the diffusion-focused expression for *μ* (Eq. 8) can be re-cast in a Michaelis-Menten form consistent with earlier ecosystems models (Eq. 9) and that our model equation set (Eq. 18a,b) is little altered from even the pioneering work of Riley (1946) and Evans & Parslow (1985). The reason for our sustained biodiversity is revealed below, but first it is useful to directly compare our findings with predictions from a previously published and very similar ecosystem model (noting here that this comparison is simply for illustrative purposes and is in no way intended to criticize the earlier model).

**Figure 5:**
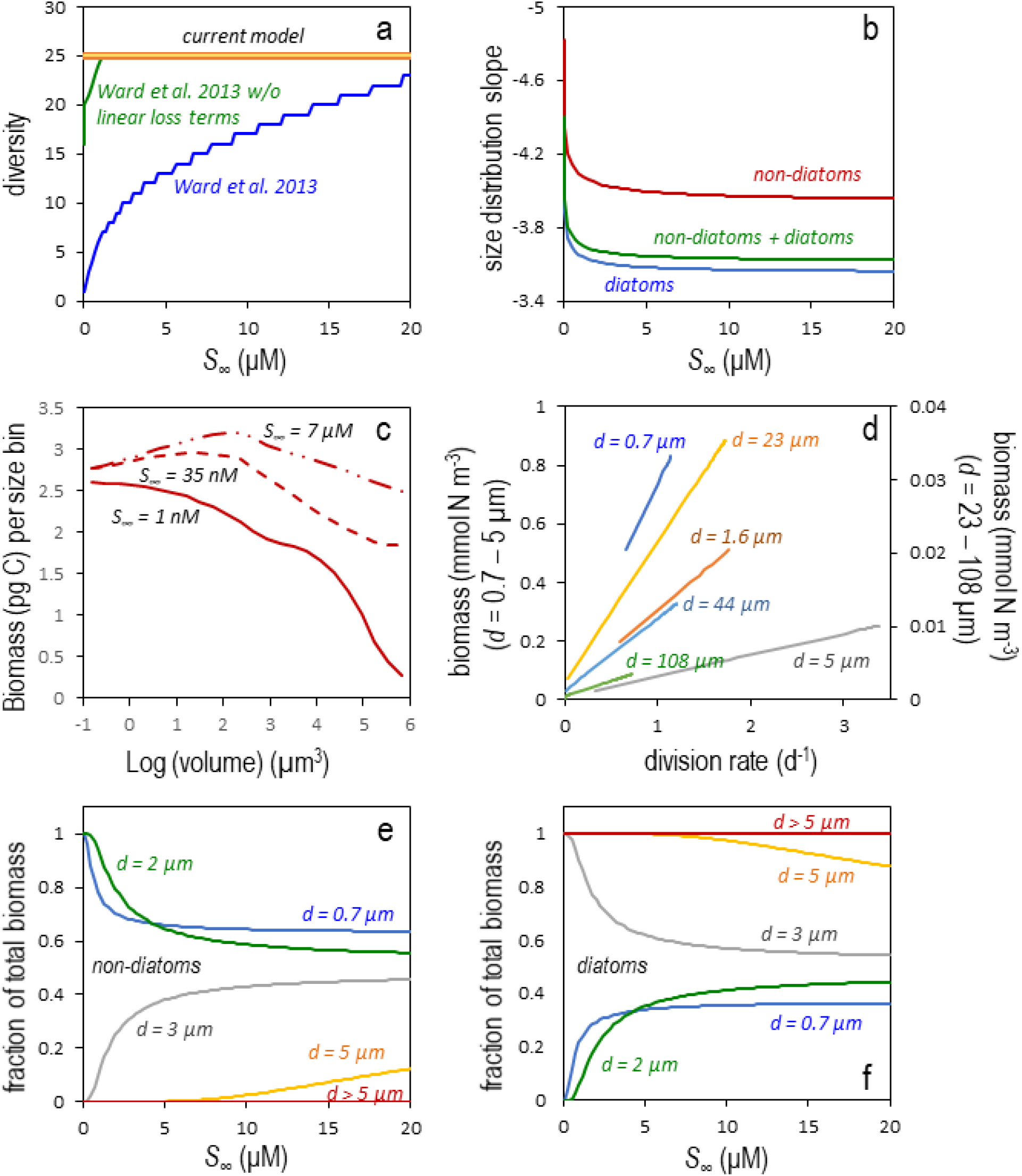
Properties of model-based steady state phytoplankton communities. (a) Phytoplankton diversity as a function of far-field nutrient concentration (*S_∞_*) for model runs initiated with 25 distinct ‘species’ (size classes). Heavy red line, thin yellow line = Non-diatom and diatom diversity for ecosystem model developed herein, respectively. Blue line = Phytoplankton diversity predicted by the Ward et al. (2013) model. Green line = Diversity predicted by Ward et al. (2013) model but with linear loss terms (*m, κ*) omitted. (b) Phytoplankton size distribution slopes (SDS) for the linear relationship between the logarithm of cell number concentration per unit length and logarithm of cell diameter as a function of *S_∞_*. Colors = Model runs for non-diatom, diatom, and mixed communities. (c) Examples of the shift in dominance from small cells to large cells as the SDS increases with increasing *S_∞_* (labeled next to line). Data are for non-diatom cell types where abundance and cell volume data values are converted to biomass following Menden-Deuer & Lessard 2000. (d) Relationships between phytoplankton division rates and biomass for cell diameters ranging from 0.7 to 108 μm. Left axis = Results for cell diameters ranging from 0.7 to 5 μm. Right axis = Results for cell diameters ranging from 23 to 108 μm. (e) Fraction of total phytoplankton biomass from the multispecies model runs that is attributable to different size classes of non-diatoms and as a function of *S_∞_*. Modeled size classes ranged from 0.7 to 135 μm, but non-diatoms only contributed significantly to total biomass at cell diameters below ~5 μm. (f) Same as in (e) except showing results for diatoms.

The Ward et al. (2013) *“idealized food-chain model”* was developed as a simplistic model to better understand the behavior of a more complex *“global food-web model”* (Ward et al. 2012). The simpler model (the code for which was kindly provided by B. Ward for developing our current model) was intended to mimic a chemostat system and thus included terms for new media input and the outflow rate of both culture media and associated phytoplankton. The Ward et al. (2013) plankton equations differ from our approach (Eq. 18a,b) in that they (1) include linear phytoplankton mortality terms (non-grazing mortality and culture outflow = 0.03 d^−1^), (2) a light/temperature-limitation term, and (3) lack a density-dependent zooplankton predation term. The Ward et al (2013) model also assumes that all small cells are non-diatoms and all large cells are diatoms. Each phytoplankton and zooplankton size class is initiated with a biomass of 10^−10^ mmol N m^−3^, which may then increase as nutrients are added to the model system. For the current comparison, we executed the Ward et al. (2013) model as originally published except with the light/temperature-limitation term removed. Predicted steady-state phytoplankton diversity from this model is shown by the blue line in figure 5a, which is essentially identical to the result presented in figure 4a of Ward et al. (2013). The model outcome is that only one or a few species (the very smallest size classes) are sustained at the lowest values of 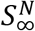 and then diversity slowly increases (additional larger species are retained) with increasing nutrient inputs.

The loss of steady-state diversity at low 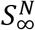 in the Ward et al. (2013) simulations is consistent with other similar models (e.g., Poulin & Franks 2010, Taniguchi et al. 2014) but it is not the consequence of resource-based competitive exclusion, as our model employs essentially an equivalent functional form for nutrient-limited phytoplankton division yet maintains all modeled species. Instead, diversity loss in the earlier model is a consequence of the non-grazing phytoplankton loss terms (linear mortality and chemostat outflow) and the initiation procedure. Specifically, the Ward et al. (2013) model is initiated with very low concentrations of phytoplankton and zooplankton in all size classes and subsequently the slow division rates of larger phytoplankton at low nutrient levels coupled with constant linear mortality (*m*) and media outflow (*κ*) rates prevent these species from accumulating to a sufficient extent that their biomass crosses even a conservative ‘extant versus extinct’ threshold (here, 0.0001% of total biomass). Indeed, at low nutrient levels, the linear mortality rate alone can exceed diffusion-limited division rates for species > 20 μm such that, even in the absence of other losses, these phytoplankton groups decrease in biomass if not repeatedly ‘restored’ to the initial 10^−10^ mmol N m^−3^. In contrast, nutrient concentrations in our model begin at a predefined 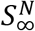, phytoplankton and zooplankton are initialized at much higher (but ecologically-relevant) biomass values, and linear phytoplankton mortality terms are excluded. The importance of non-grazing mortality as an agent for exclusion in earlier models (Poulin & Franks 2010, Ward et al. 2013, Taniguchi et al. 2014) is revealed when the model is re-run with the *m* and *κ* terms removed, which results in nearly full diversity being sustained at even the lowest 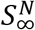 (Fig. 5a, green line).

In addition to sustaining size diversity, our ‘baseline’ model predicts phytoplankton abundances in the smallest size bin (equivalent to *Prochlorococcus*) of ~10^5^ cells ml^−1^ and larger species that follow a phytoplankton size distribution slope that tilts upward (i.e., become less negative) as 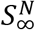 increases from 1 nM and 20 μM (Fig. 5b). The model range for the size distribution slope and its behavior with 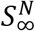 is broadly consistent with field observations (e.g., Sheldon 1972, Huete-Ortega et al. 2012, Marañón 2015, Behrenfeld et al. 2021a) and corresponds to biomass dominance shifting between picophytoplankton and microphytoplankton (Fig. 5c). These shifts in community structure are driven by size-dependent potentials for diffusion-driven increases in *μ* as 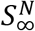 increases (Fig. 5d), noting here that these relationships are not 1:1 with biomass and reflect predator-prey balances between division and loss rates at equilibrium (Behrenfeld & Boss 2014, 2018).

Results presented here indicate that our ‘baseline’ model yields properties of phytoplankton communities reflective of field observations and provide an explanation for the loss of diversity in previous modeling studies. While it is beyond the scope of the current study to fully explore other model parameterizations or constructs, it is hard to resist the temptation to try at least one modification. In particular, what happens when diatoms and non-diatoms are modeled together across all size classes? To answer this question, we modified the baseline model such that each of the 25 phytoplankton size classes had a diatom and non-diatom representative (each initiated with 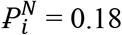 mmol N m^−3^) and we assumed that zooplankton grazers in each size class had no preference for phytoplankton prey types. All other aspects of the model were unaltered. Steady-state size distribution slope solutions for this multispecies model are shown as a function of 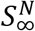 in figure 5b (green line), while the fractional contributions of non-diatoms and diatoms to total phytoplankton biomass in each size class are shown in figures 5e and 5f, respectively. The multispecies size distribution slopes (Fig. 5b, green line) reflect shifts in community composition, with non-diatoms dominating at low 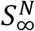 and diatoms dominating at high 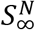. The reason for this shift is that swimming by non-diatoms proffers a minor advantage over diatom in terms of diffusive nutrient flux at the smallest cell sizes. In contrast, sinking and cell vacuolation (accounted for in the Menden-Deuer & Lessard 2000 relationship) slightly improve nutrient acquisition relative to requirements in large diatoms compared to swimming in unvacuolated non-diatoms. While these differences in diffusive flux are small, they are associated with slight changes in division rate that, when played out over time, result in resounding size-dependent shifts in species dominance (Fig. 5e,f), a reflection of the ‘trophic exclusion’ principle discussed in Behrenfeld et al. (2021b).

## Synthesis

The realm of the phytoplankton can be non-intuitive to large-bodied terrestrial organisms such as ourselves. The number of individual phytoplankton in a waterbody may be astronomical, but from a body-length perspective they are distantly spaced. The tempo of phytoplankton division and death is unfamiliar to us, yet it is of fundamental importance to community structuring and succession. A limiting nutrient may be only a short distance from a cell, but accessing this resource can be challenging because most phytoplankton have a limited capacity to move relative to the water molecules surrounding them. In three previous studies (Behrenfeld et al. 2021a,b,c), we grappled with developing an understanding of diversity and succession in this foreign world experienced by phytoplankton, and it seemed to us then that a key in doing so would be to explicitly account for the discreteness of individual cells. With this in mind, we undertook the current investigation to model phytoplankton populations from the ‘perspective’ of a cell, albeit not by modeling individual cells *per se*. Within this perspective, phytoplankton are immersed in a medium where far-field limiting nutrient concentrations are a constant (i.e., assimilation and remineralization rates are balanced), there is no direct resource competition between neighboring cells, and performance of a given species (cell size) is defined by nutrient diffusion across a boundary layer and a physiological optimization strategy that, at sufficient resource supply, supports an evolutionarily-selected maximum growth rate. Our hope was that this approach might help illuminate why extreme competitive exclusion appears to be a common behavior in contemporary ecosystem models.

One of our initial concerns with constraining phytoplankton growth strictly through diffusion limitation was that resultant division rates would unrealistically vary inversely with cell diameter squared. It was therefore satisfying when size distributions in division rate emerged from our semi-empirical model that varied by even less than the surface:volume ratio for nutrient concentrations representative of nearly all natural waters (Fig. 3). The reason for this outcome is that, over much of the phytoplankton size domain, realized division rate is governed more by the optimization of cellular machinery than diffusional flux, even at the lowest nutrient levels. In other words, smaller cells are operating in the slowly-saturating region of their *μ-S_∞_* relationships (Fig. 2d) where diffusion potential exceeds utilization. It is noteworthy, here, that our modeled division rates essentially represent an upper limit on growth (if not supplemented by other nutrient sources; e.g., mixotrophy) at low nutrient concentrations that, apparently, some species have evolved to take full advantage of (Fig. 4a) and others have not (Fig. 4b).

In Behrenfeld et al. (2021a,b), we expressed a concern that contemporary ecosystem models encompass an unrealistic degree of resource-based competition between phytoplankton classes, a view earlier shared in Siegel (1998). Our mistake was in envisioning that the modeling of phytoplankton groups simply as integrated elemental stocks was equivalent to treating them as diffuse overlapping fields (i.e., ‘fluid variables’) where direct competition is continuous. By formulating growth herein as a function of boundary layer diffusion, it seemed to us that a model could be developed that is completely compatible with phytoplankton being discrete entities in a competition-neutral resource landscape. However, in the process of developing this model, we realized that our diffusion-focused equation could be transformed into a mathematically-equivalent Michaelis-Menten form consistent with earlier model formulations. This insight (at least for us) made it clear that direct resource competition is not inherently implied by treating phytoplankton biomass as an integrated elemental mass and that observed extinctions of phytoplankton size classes in ecosystem models cannot simply be interpreted as competitive exclusion by smaller cells with higher nutrient ‘affinities’. Indeed, we propose that the common association of nutrient half-saturation values 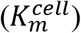 with ‘affinity’ has misguided earlier interpretations and that 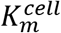 is better viewed, going forward, as simply an emergent trait reflecting a size-dependent diffusion constraint and evolved strategies for up-regulating cellular capacities with increasing resource availability. In the discrete world of the phytoplankton, there is rather little a cell can do beyond sinking and swimming to enhance diffusive potential and, even if it could, it generally would have little impact on the far-field nutrient environment experienced by neighbors.

With the above realizations, it became even more confusing as to why ecosystem models generally predict low phytoplankton biodiversity in nutrient impoverished regions, a prediction opposite that of observations (Ibarbalz et al. 2019). By employing our diffusion-governed growth equations in a phytoplankton-zooplankton equation set, we find that the entire biodiversity included in our ‘baseline’ model at initiation is sustained within the emergent steady-state populations for all far-field nutrient concentrations. Instead of nutrient competition being the cause of exclusions in models, we find that these earlier-reported species losses are largely attributable to very low model initiation concentrations and, more importantly, the inclusion of a linear phytoplankton mortality term that is described in a manner, as we argue below, that is difficult to justify.

As commented at the beginning of the previous section, ocean ecosystem modeling is a bit of an art form and, at this, we must confess we (particularly, M.J.B.) are rather rudimentary artists. There are many facets of plankton ecology that our simple ecosystem model fails to capture. For example, our model does not account for competition between individuals when relative motions cause boundary layers to transiently overlap and we don’t explicitly account for phytoplankton losses to viral lysis (although, if this processes is density-dependent, it might be envisioned as included in the grazing term of equation 18a). Additionally, our model fails to address the importance of mixotrophy in phytoplankton growth rates (e.g., Flynn et al. 2018, Ward et al. 2011), we do not account for the roles of selective feeding or grazer defense strategies, and we only consider the condition of uniform nitrogen limitation, ignoring the effects of nutrient patchiness (Mitchell et al 2013), potential unique aspects of phosphate, iron, or light limitation, and potential advantages of hosting endosymbionts (Caputo et al. 2019, Decelle et al. 2012, 2019). Despite these omissions, we believe the fundamental conclusions state above are robust.

An important question yet to address that emerges from our study is how to capture non-grazing phytoplankton losses in future modeling? The fact is that phytoplankton do die in nature from processes other than being eaten. For example, stress can lead to programmed cell death (Bidle 2016), viral lysis can behave in a manner that is not density-dependent (Knowles et al. 2020), and other forms of disease and life cycle transitions can result in phytoplankton loss (Brussaard & Riegman 1998, Choi et al. 2017). However, without a mechanistic understanding on how to predict when, where, and to what extent these mortality processes occur, it is difficult to accurately represent them in models (Taniguchi et al. 2014). Certainly, accounting for these loss by treating them as a constant, albeit low, daily loss rate is unrealistic and here we have shown that this approach can actually have catastrophic impacts on modeled community diversity. Perhaps, it may be more prudent to simply omit such linear mortality terms in models until we have developed a fuller understanding of their drivers.

Throughout this manuscript, we have emphasized the role that spatial distancing between phytoplankton cells plays in diminishing potentials for direct resource-based competition. It is important to understand, however, that we are not suggesting that the plankton world is free of competition. Rather, we envision it as a landscape of extreme competition, but one that is largely not based on resource acquisition. Instead, this competition plays out through the interactions of predators and prey. The average life expectancy of an individual phytoplankton in the global ocean is on the order of a day to weeks (Behrenfeld & Falkowski 1997). Under this rapid tempo of turnover, minor differences in fitness between species result in selection of a finely tuned biodiversity, a process we earlier referred to as the ‘trophic exclusion principal’ (Behrenfeld et al. 2021b). An example of this principal is illustrated in figures 5e and 5f, where minor differences in division rates between phytoplankton groups led to a size-dependent selection for non-diatoms or diatoms. If in this scenario the diffusion-based differences in division rate are countered by reduced grazing in small diatoms and mixotrophy in larger non-diatoms, then one can imagine how a re-parameterized model could yield a sustained coexistence of both phytoplankton groups across all size classes and far-field nutrient concentrations. The key point here is that ‘fitness’ is defined by any adaptation that allows persistence of a given species within a community, not simply acquisition of growth limiting resources. Importantly, the time-scale over which fitness is selected can be very long (years), causing shorter-term species-specific advantages to be averaged out and enabling greater sustained diversity within functional size classes (Behrenfeld et al. 2021b).

## Acknowledgements

We thank Ben Ward for graciously providing computer code for his idealized food chain model, Kimberly Halsey, Allen Milligan, and Toby Westberry for data support and helpful discussions. MJB was supported under NASA grants NNX15AF30G (NAAMES), 80NSS17K0568 (EXPORTS), and 80NSSC22K0358. KB was supported by NASA grant 80NSSC20K0970. EB and LKB by NASA grant 80NSSC21K0783. PG was supported by NASA PACE Science Team grant 80NSSC20M0202.

## Footnotes

* For this size distribution, the differential number concentration, *N*(*d*) [cells ml^−3^ μm^−1^] is given by 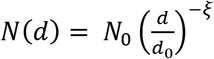, which when integrated between *d_min_* and *d_max_* gives equation 2.

† For these modeled populations and assuming *Prochlorococcus* to span a size range of 0.6 to 0.8 μm diameter, the value for *N*_0_ ranges from 2.9×10^−5^ to 1.8×10^−4^ cells ml^−1^ μm^−1^ and can be calculated as: 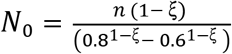

‡ Note that an *I_r_* term would also need to be added to equation 5 if *P* had been measured as net oxygen production in Fig. 2a, rather than ^14^C uptake

§ The *I_r_* term appears in equation 6 because catabolism of carbon products ultimately results in the respiratory production of CO_2_, which can isotropically diffuse through the cell and across the outer membrane. In contrast, catabolism of nutrient-containing molecules generally does not proceed to inorganic forms, but rather to intermediate or charged forms that are readily re-assimilated into new products and unlikely to be lost from the cell in significant quantities. For example, nitrogen-containing molecules are degraded to simple amino acids, ammonium, or urea. Accordingly, a term analogous to *I_r_* is not included in equation 8.

** Note that the ‘10,000’ factors in equations 12a and 12b are for units conversion. The original relationships were developed with diameter in units of cm and velocities in cm s^−1^. In the current manuscript, diameters have units of μm and velocities have units of μm s^−1^.

†† The chosen threshold defining whether a modeled phytoplankton species is extant or extinct is intended to be conservative. To put this threshold in perspective, we expect large phytoplankton to be very rare (but not extinct) in oligotrophic waters. If we assume that *Prochlorococcus* has a cell abundance of ~10^5^ cell ml^−1^ and that the phytoplankton size distribution has a steep slope of −5.3, then the abundance 135 μm cells (the largest size class represented in our model) would be ~1 cell per 10 m^3^ and its contribution to total phytoplankton biomass would be ~0.0008%, which exceeds our 0.0001% threshold defining whether a modeled species is extant or extinct.

## Notes

### Competing Interest Statement

The authors have declared no competing interest.

